# Detection of Motor-Evoked Potentials Below the Noise Floor: Rethinking the Motor Threshold

**DOI:** 10.1101/2021.12.10.472126

**Authors:** Zhongxi Li, Angel V. Peterchev, John C. Rothwell, Stefan M. Goetz

## Abstract

**Background:** Motor-evoked potentials (MEP) are one of the most prominent responses to brain stimulation, such as supra-threshold transcranial magnetic stimulation (TMS) and electrical stimulation. Understanding of the neurophysiology and the determination of the lowest stimulation strength that evokes responses requires the detection of even smaller responses, e.g., from single motor units. However, available detection and quantization methods suffer from a large noise floor.

**Objective:** This paper develops a detection method that extracts MEPs hidden below the noise floor. With this method, we aim to estimate excitatory activations of the corticospinal pathways well below the conventional detection level.

**Methods:** The presented MEP detection method presents a self-learning matched-filter approach for improved robustness against noise. The filter is adaptively generated per subject through iterative learning. For responses that are reliably detected by conventional detection, the new approach is fully compatible with established peak-to-peak readings and provides the same results but extends the dynamic range below the conventional noise floor.

**Results:** In contrast to the conventional peak-to-peak measure, the proposed method increases the signal-to-noise ratio by more than a factor of 5. The first detectable responses appear to be substantially lower than the conventional threshold definition of 50 μV median peak-to-peak amplitude.

**Conclusion:** The proposed method shows that stimuli well below the conventional 50 μV threshold definition can consistently and repeatably evoke muscular responses and thus activate excitable neuron populations in the brain. As a consequence, the IO curve is extended at the lower end, and the noise cut-off is shifted. Importantly, the IO curve extends so far that the 50 μV point turns out to be closer to the center of the logarithmic sigmoid curve rather than close to the first detectable responses. The underlying method is applicable to a wide range of evoked potentials and other biosignals, such as in electroencephalography.

## Introduction

Motor-evoked potentials (MEPs) are an important response phenomenon in brain stimulation, such as transcranial magnetic stimulation (TMS). If motoneurons in the human primary motor cortex are activated directly or indirectly, they respond with action potentials, which are transmitted down the spinal cord to the lower motoneurons [1]. The lower motoneurons route the signals to the muscles, where they can be detected as MEP waves through electromyography (EMG). MEPs provide one of the few directly observable responses of the brain to stimuli. Furthermore, MEPs are a key to understanding the biophysical and neurophysiological mechanisms of brain stimulation, to establishing safety limits, to matching different devices or coils, and to achieving individual dosing [2-5]. The so-called motor threshold is defined based on MEPs as a low stimulation amplitude that generates the first reliably detectable responses and is the key safety as well as dosage reference level [6].

The response amplitudes of several MEPs graphed over the corresponding stimulation strengths forms an s-shaped curve, often named as input–output (IO) or recruitment curve [7-10]. For high stimulation strengths, the responses saturate and form the high-side plateau of the sigmoidal IO curve around several millivolts, dependent on the specific muscle [11]. If the stimulation strength is reduced, the response amplitude decreases monotonically until the responses appear to cease entirely and fall below a low-side plateau or noise floor that is formed by endogenous activity, biosignals from unrelated neural and muscular sources, and recording noise [10]. Thus, the dynamic range of MEP amplitudes can exceed a factor of 1000 [12].

Typically, the noise floor forming the low-side plateau for a resting motor system of a conscious subject, e.g., for TMS, can reach above 10 μV peak-to-peak [13]. The motor threshold was defined to be slightly above this plateau at 50 μV peak-to-peak median response amplitude as a compromise between detecting subtle responses and sufficient detection robustness [12]. Such threshold definitions assume that they are close to the smallest occurring evoked activity. The noise floor, however, is not a purely technical property of the used amplifier, but—for modern amplifiers—widely depends on externalities, such as the electrode and source impedance, body temperature, and biosignals from unrelated sources [4]. Thus, simple technical advances of amplifiers do not solve the problem. According to the established interpretation, smaller responses causally evoked by stimulation of motor efferences might either not exist or could not be detected for physical reasons.

Particularly for neuromodulatory interactions, e.g., repetitive TMS with certain pulse rhythms or patterns, the threshold is physiologically important beyond amplitude individualization and safety [14]. Many protocols, particularly inhibitory 1 Hz as well as patterned protocols such as quadripulse and theta-burst stimulation, use sub-threshold pulses, which is often below 90%, and sometimes below 70% of motor threshold [15-19]. In fact, the pulse amplitude during the modulatory intervention can determine if the intervention is inhibitory, excitatory, or effectless [20-23]. Such definitions may leave the impression that there could be a rather binary threshold, which may have a stochastic nature or blurring due to the high trial-to-trial variability but still demarcate a transition at the threshold or very close to it.

Early research on inhibition also considered that fatigue mechanisms both in the central and the peripheral nervous system as well as in muscles might be involved [24, 25]. Such considerations may encourage rather using subthreshold repetitive protocols and avoiding substantial activation of muscle units. More recent paradigms might want to rule out motor activation and at most aim at some inhibitory interneuron activation. However, do subthreshold stimuli really not activate corticospinal connections and how weak does a stimulus have to be to avoid activation?

So-called active stimulation into a pre-activated motor system is used to find smaller effects of single TMS pulses [26]. However, it is not obviously clear if such active pulses actually immediately depolarize neurons or only modulate endogenous signals. First approaches for looking below the noise floor and finding MEPs averaged the responses to many individual stimuli, as a solution that is also typical for other subtle biosignals, such as in electroencephalography (EEG) [27]. Although these experiments hinted that there is evoked motor activity well below the motor threshold, averaging does not solve the problem. Averaging of responses to several different stimulation strengths or even of an entire IO curve would require many trials and lead to unacceptably long session durations. More importantly, however, MEPs are inherently highly variable, which reflects endogenous signaling and modulation of the neuronal target [13, 28]. Averaging extinguishes this variability and the associated information. Furthermore, the variability is not Gaussian but has a highly skewed distribution. Consequently, averaging does not result in an exemplary representation of the many variable and noisy individual MEPs, but is dominated by outliers [29]. Thus, within a large series of repetitions, very few rare and maybe even spontaneous potentials with the right timing dominate and determine the averaged response.

We propose an adaptive matched-filter method for reliably detecting the MEPs below the noise floor. The method treats each MEP as a composition of multiple motor unit action potentials (MUAPs) that are identical in shape but with different latencies and amplitudes. The latencies and amplitudes of the MUAP components are estimated for each MEP with maximum likelihood, while the MUAP shape itself is learnt from previous responses of the same subject. The approach achieves a high signal-to-noise ratio (SNR) through the extraction of the shared MUAP pattern across many responses and therefore shares some characteristics with averaging but translates them to the individual MEPs.

In contrast to simple time-synchronized averaging approaches, the proposed method has several advantages. First, it does not require repetitions at the same stimulation strength but merges information from all applied valid stimuli with a wide range of stimulation strengths. Second, it does not introduce bias due to the non-Gaussian distribution of MEPs and is resilient to infrequent spontaneous potentials in an averaged ensemble [30, 31]. Third, it provides a quantitative response amplitude estimate, such as equivalent peak-to-peak voltage amplitude, for each MEP so that the physiological variability is maintained for subsequent analysis steps and is therefore compatible with all known MEP-based methods such as thresholding or IO curves [32-34]. Finally, it reveals the distribution of the MUAP population over time per MEP recording, providing detailed latency information that is insensitive to measurement noise.

## Material and methods

Conventionally, the amplitude of MEPs is quantified by their peak-to-peak voltage [35]. While simple, this approach is relatively insensitive to weak MEP waves but highly susceptible to noise. The peak-to-peak measurement is likely to pick outliers of distributions and converts interferences and noises, such as additive Gaussian noise, into highly skewed extreme-value distributions [12, 36]. Strictly, those extreme-value distributions have to be considered for correct statistical analysis, but practically rarely are.

Previous studies suggested using the area under the MEP curve instead of the peak-to-peak reading for greater accuracy and higher tolerance to noises [13, 37, 38]. However, several disadvantages exist: the reading depends on the amplifier filter properties, such as the high-pass filter cutoff; it requires a definition of start and end of an MEP, which varies substantially at weak stimulation strengths; it provides impractical units (volt-seconds) and is incompatible with traditional peak-to-peak quantification as well as established safety rules [39]. Most importantly, the MEP area extraction either needs prior information about beginning, zero-crossing, and end or has to rectify the signal, which introduces bias due to converting Gaussian noise into a one-sided distribution and therefore does not substantially increase the sensitivity. As a consequence, the MEP area is rarely used in practice.

Our earlier work proposed a matched-filter method to extract the EMG amplitude [4]. This method can detect EMG signals that are substantially smaller than the noise floor. However, the method still neglects the variable latency of the response MEP. Fixing the differential latency enabled higher sensitivity with fewer samples by reducing the degrees of freedom. Although variable latency could be accounted for by time-shifting the matched filter, this approach could be inconsistent if the time shift is not inherently estimated and inaccurate for responses with very different latencies.

### Assumptions

We assume that each MEP measurement comprises a volley of multiple motor unit (compound) action potentials (MUAPs) that share the same shape but with different latencies and amplitudes. Specifically,

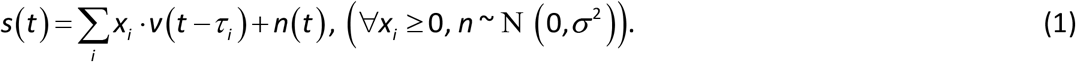

where *s*(*t*) is the MEP recording, *v* the MUAP, *X*_*i*_ the amplitudes of MUAPs at different time, *τ*_*i*_the latencies, and *n* the Gaussian noise, while subscript *i* enumerates the different possible latencies. For causal responses to the stimulus, we restrict the response to *X*_*i*_ ≥ 0 for all possible latencies *i*. The concept of the MEP composition is visualized in Fig. 1. Thus, with known MUAP shape, we should ideally extract the population of MUAPs over time (*X*_*i*_, *τ*_*i*_) for each MEP waveform. Per MEP, the MUAP populations reflect the amplitude of the MEP response.

**Fig. 1.**
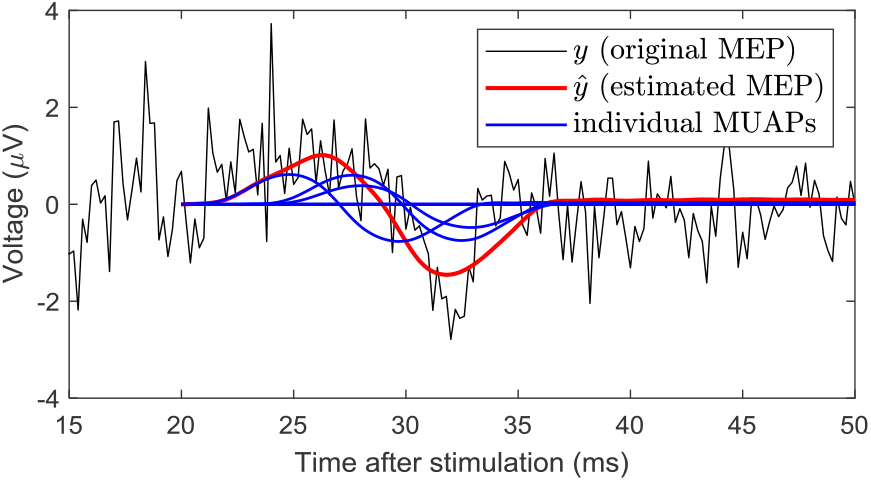
An example of MUAP decomposition for a weak MEP response.

### Decomposition and evaluation

If the MUAP shape *v* is known, the weights *X* can be estimated by a maximum-likelihood approach:

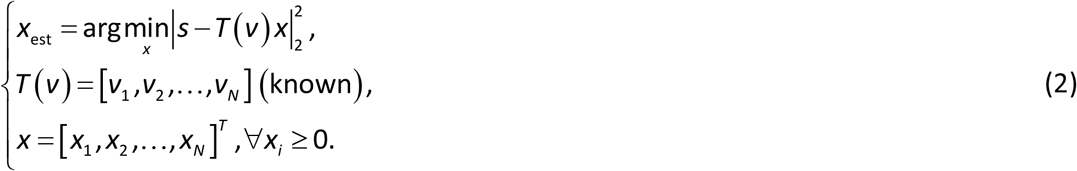

In this setup, we enumerate the possible delays of MUAPs by *i*. The actual time resolution is ms per the sampling rate of the recording device. Matrix *T*(*v*) gathers all time-shifted MUAP templates *v*_*i*_ in a way similar to a Toeplitz matrix except with zero-padding. Each column of *T*(*v*) matches *i*, and the number of columns reflects the detection time window. In this study, we set the detection window to be 20 ms ≤ *t* ≤ 44 ms, which parsimoniously covers the full dynamics of the MEP while avoiding the TMS artifacts. Vector *X* gathers the unknown weights with the dimension matching that of *T*(*v*).

Equation (2) is a typical quadratic programming problem with linear constraints, which can be solved efficiently. In this paper, we use the *quadprog* solver in MATLAB. Equation (2) resembles matched filter methods in that the best amplitude estimation at the presence of Gaussian noise solves a least-squares problem. The difference is that the proposed method additionally considers unknown latencies and ensures non-negative weights (*X*_*i*_). Limiting the amplitudes to non-negative values corresponds to our assumption that each MEP is a sum of actual MUAPs and, henceforth, allows physiological interpretations of the decomposition results.

After the decomposition, the noise-free MEP is estimated by

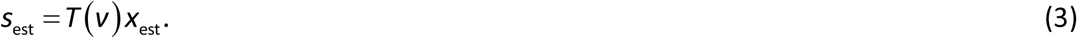

Both *S*_est_ and *X*_est_ can give noise-free quantification of the strength of an MEP response. For example, an element-wise summation of the amplitudes Σ *x*_est,*i*_ describes the response strength since it reflects the total recruitment population. Additionally, the individual entries of *X*_est_ provide detailed insights of the time delay. Since *s*_est_ is practically the MEP without noise, one can of course use max(*S*_est_) − min(*S*_est_) to provide a noise-free estimation of the peak-to-peak voltage Vpp as an MEP reading, which is commonly used by the community.

### Recovery of the MUAP shape

If the MUAP weights *X* are known, the best estimate of the MUAP shape *v* follows the minimization

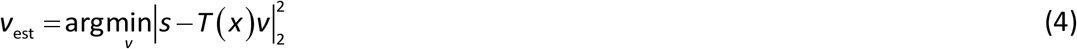

where matrix *T*(*X*) organizes the unknown weights *X* in the same way as *T*(*v*) organizes *v* in (2). As such, *T*(*X*) *v* = *T*(*v*) *X*, both attempting to approximate the ground truth *s*. We choose different formalisms for Equations (2) and (4) in order to expose different unknowns as a linear argument, making both problems solvable in the same manner. Such arrangement is mathematically possible because the estimated MEP *T*(*v*) *X* is a bilinear function of *v* and *X*.

We combine (2) and (4) to estimate the MUAP shape *v* and weights *X* iteratively:

1. Initialize MUAP shape *v*;
2. Feed *v* to Eq. (2) to update *X*;
3. Feed *X* to Eq. (4) to update *v*;
4. Repeat steps 2–3 until *v* converges.

In this paper, we initialize the MUAP shape *v* as the stimulus-synchronized average MEP waveform per subject. Over the iterations, the MUAP shapes deform slightly and stabilize after 4−5 iterations. The MUAP shape develops higher-frequency features and a briefer shape overall after sufficient numbers of iterations (see Fig. 1), which is expected as the MEP is a sum of time-shifted MUAPs and therefore mathematically a low-pass filtered product of the MUAP shape in this formalism. Although the final MUAPs still resemble the MEP waveform (Fig. 2), the approximation error, measured as mean(|*s* − *T*(*x*)*v*|_2_^2^) per subject, is reduced by 40%−50%. The iterative algorithm demonstrated robustness since similar results can be obtained from completely random initializations. However, we stuck to the heuristic average-MEP initializations for the analysis here to avoid local minima.

**Fig. 2.**
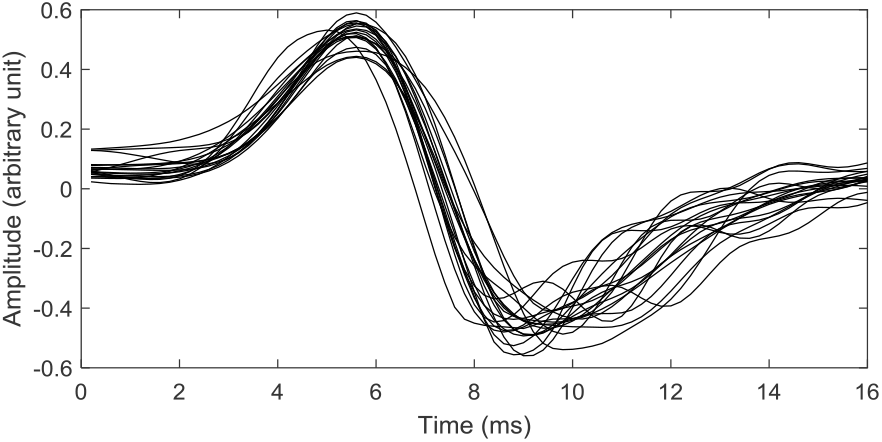
Learned MUAP shapes from 21 subjects (one MUAP per subject).

### Sensitivity to noise

To benchmark the sensitivity of the proposed method, we generate artificial MEP waveforms using a learned MUAP shifted by random weights *x* as *ground truth*. Gaussian noise is additionally introduced with controlled variance to effect different signal-to-noise-ratios (SNRs). Two detection methods are compared: the proposed method and the conventional peak-to-peak reading. Both methods attempt to evaluate the latency and amplitude of the underlying true MEP.

The evaluation results are shown in Fig. 3 where Row (a) shows an overview of the Vpp estimation error under different SNRs, Rows (b–c) detail the estimation errors of Vpp and latency at selected SNRs, and Row (d) visualizes waveforms for three SNRs. The proposed method has substantial advantages in terms of both Vpp and latency estimation. In particular, the proposed method provides an unbiased estimate of Vpp while the conventional method is susceptible to noise and systematically overestimates the Vpp of weak MEPs.

**Fig. 3.**
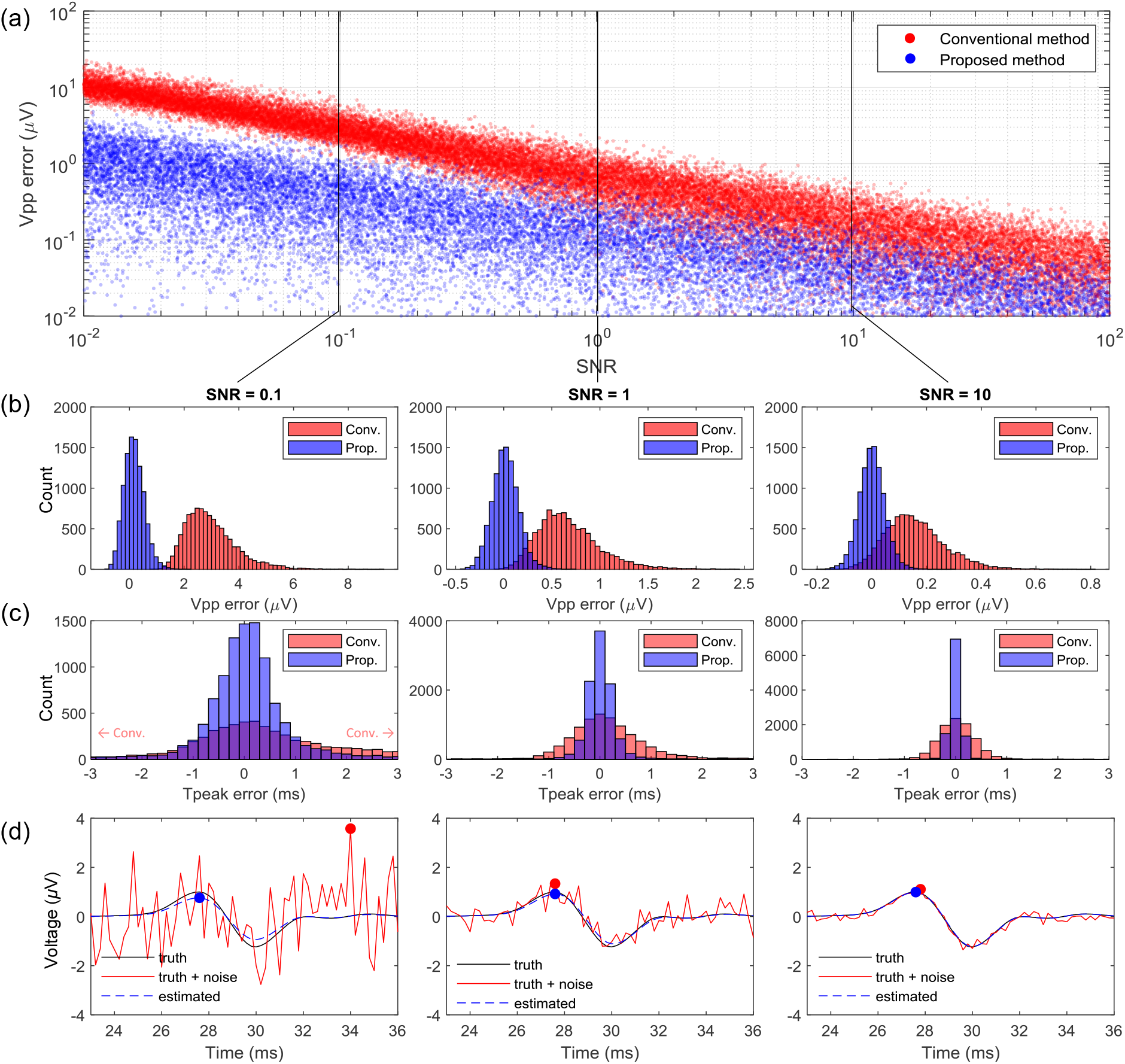
Sensitivity analysis of conventional peak-to-peak detection vs. the proposed adaptive learning estimation method. Both the ground truth MEP and the noise are simulated. Row (a): Vpp estimation error for different signal-to-noise ratios (SNRs). Rows (b–c): Histograms of the Vpp estimation error and the peak time error for SNR = 0.1, 1, and 10. Row (d): Representative waveforms sampled from SNR = 0.1, 1, and 10. Each column of Rows (b–d) has the same SNR.

Fig. 4 presents a *sanity check* of false positives, where both detection methods are applied to pure Gaussian noise. The detected MEPs are interpreted as *false alarm* and should ideally be zero. Again, the proposed method outperforms the conventional one with a large margin.

**Fig. 4.**
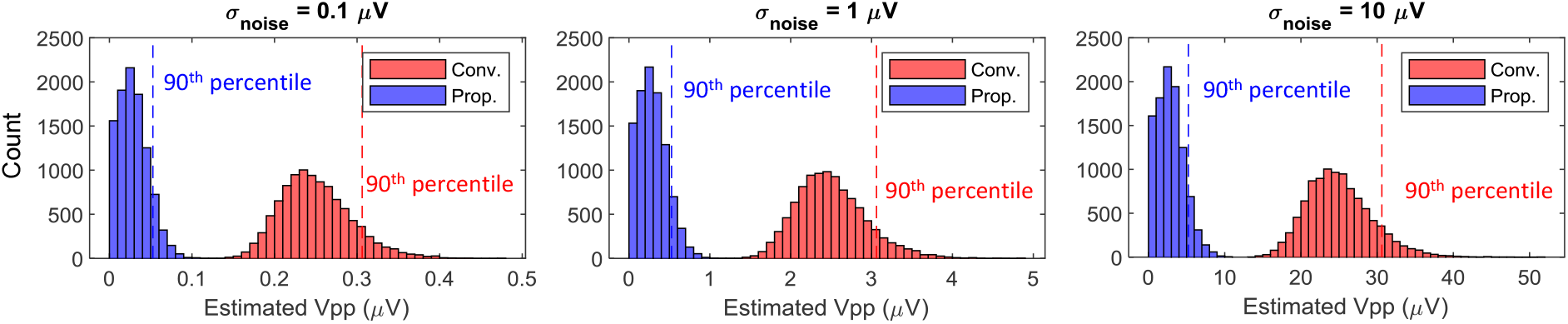
Detection results on Gaussian noise (no MEP contained). An ideal detection method should yield Vpp = 0 due to the absence of MEPs.

When processing real data, we generate the same performance chart as Row (b) of Fig. 3 but refer to the actual standard deviation of the noise, σ_noise_. The performance chart is used to infer a 90th percentile noise floor (in units of μV) in IO plots.

## Results and discussion

We evaluated the method experimentally and combined the data acquisition with an earlier study [40]. The procedure was approved by the Duke University Institutional Review Board. The subjects underwent single-pulse TMS over the representation of the first dorsal interosseous (FDI) muscle in the left primary motor cortex. A focal figure-of-eight coil (MagVenture B65, Farum, Denmark) was positioned approximately 45° to the interhemispheric fissure to align the induced electric field perpendicularly with the hand-knob gyrus. The coil position with maximum MEP output amplitude in the FDI muscle was identified manually before the procedure. A robotic coil holder maintained the position of the coil throughout the session and compensated subject movements, while subjects were instructed to sit as still as possible [41, 42]. MEPs were recorded and sampled synchronously to the TMS pulse trigger through surface Ag/AgCl electrodes and an MEP amplifier (K800 with SX230FW pre-amplifier, Biometrics Ltd., Gwent, UK) at 5 kHz and 16 bit. Recordings with activity of more than 40 μV within a window of 200 ms before the TMS pulse were marked as facilitated and excluded from the analysis. The individual pulses were at least 8 s apart and inter-pulse timing was randomized to not follow the subject’s expectation or pick up any regular excitability oscillations. We recorded IO curves with randomized stimulation amplitudes for each subject.

We performed the above routine for 21 subjects. The learned MUAP shapes vary by subject but are similar overall (Fig. 2).

### IO curves and how weak can stimuli be to still evoke excitatory activations

Fig. 5 displays IO curves extracted from the data with conventional peak-to-peak MEP amplitude measurement and the proposed matched-filter estimator. The 90th noise floor overlays in each IO plot mark the background activities. The background activities are gauged from the pre-stimulation MEP recordings, using the corresponding extraction methods as if they are actual post-stimulation recordings. The times of MEP peaks are extracted as follows: for the conventional method, the peak times are obtained by directly finding the peak of the raw recording; for the proposed method, the peak time is obtained from the reconstructed signal *T*(*v*) *x*.

**Fig. 5.**
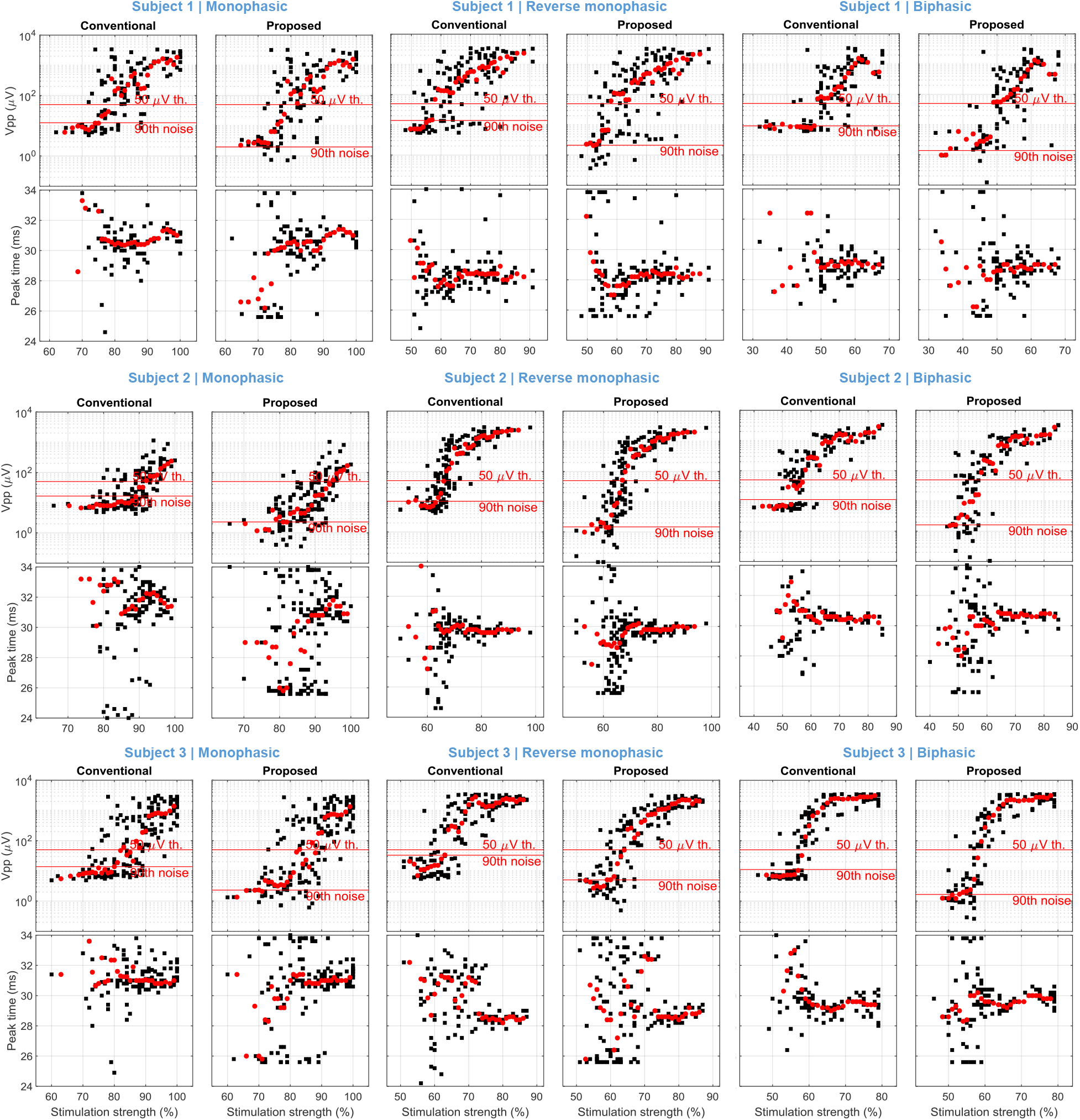
IO curves and peak EMG times (black) with moving-window median (red) in three example subjects for three different TMS pulse configurations (monophasic, reverse monophasic, and biphasic). Two types of detection methods are compared. “90th noise” refers to the background activity recorded shortly before the stimulation, using the same extraction method as if they are post-stimulation MEPs. The current direction is normal unless otherwise denoted in the titles.

For weaker stimuli, the conventional method collapses at around the 90th percentile of background activity and fails to distinguish MEP responses therein. Conventional peak-to-peak measurement leads to a lower-side plateau above 5 μV. The proposed estimator, in contrast, is less sensitive to noise as confirmed by the previous section so that responses can be detected and quantified notably below the 5 μV technical noise floor. These weak responses—which were usually considered noise—actually form a noticeable trend that reflects the expected positive stimulation-strength dependency. The trend monotonically decreases until reaching near the 90th background activity baseline (the “90th noise” marks of Fig. 5). These new baselines are around ^1^/_5_ (−14 dB) of their conventional counterparts.

As an important observation, the commonly used 50 μV threshold, i.e., the stimulation amplitude that leads to a median response of 50 μV peak-to-peak, no longer marks the onset of first detectable MEPs, but is almost in the center of the sigmoid (see Fig. 5). Thus, it turns out that the point that once was considered a stimulation strength at a level where first detectable responses occur is rather in the middle of the dynamic range of MEPs. The above observations are substantiated by further data shown in Fig. 5. The monotonic trend of weak MEP amplitudes (above the noise floor but below the 50 μV point) is ubiquitous, especially when more measurements are available.

## Conclusions

We presented a method to detect MEPs in response to TMS. The method is notably more sensitive than conventional peak-to-peak metrics so that responses below the noise floor could be detected. Our analysis indicated that the detection sensitivity could be increased by 14 dB.

In addition to a detection of even smaller MEPs in response to weak stimuli due to the increased sensitivity, the method can also detect the already established dynamic range of the IO curve and provide the same reading in peak-to-peak voltage with lower distortion and higher accuracy. This peak-to-peak voltage of the ideal, noise-free MEP tends to turn out smaller in amplitude, whereas the conventional peak-to-peak detection is highly sensitive to noise, which introduces asymmetric bias and therefore systematically overestimates MEPs.

The formalism is based on matched- or optimum-filter estimation, which is a well-established maximum-likelihood-grade detector in communication systems. Since, in contrast to communication systems, the optimum filter is not known and is individual, the method learns it adaptively on the go. The MEP detection filter or pattern is iteratively improved, and previous measurements can be continuously reappraised with the improved filter.

The method uncovers that stimuli well below the conventional 50 μV threshold definition can consistently and repeatably evoke muscular responses and thus activate excitable neuron populations in the brain. As a consequence, the IO curve is extended at the lower end, and the noise cut-off is shifted. Importantly, the IO curve extends so far that the 50 μV point turns out to be closer to the center of the logarithmic sigmoid rather than close to the first detectable responses.

Finally, the presented method is applicable to other forms of brain or peripheral stimulation as well as potentially to other biosignals representing evoked responses.

## Availability

The method used for the analysis and instructions are available at https://github.com/zlduke/MEP-decomposition to the community for free use and further development.

